## Supplementary figures and images for "Transcriptome association studies of neuropsychiatric traits in African Americans implicate *PRMT7* in schizophrenia"

### Supplemental Figure 1

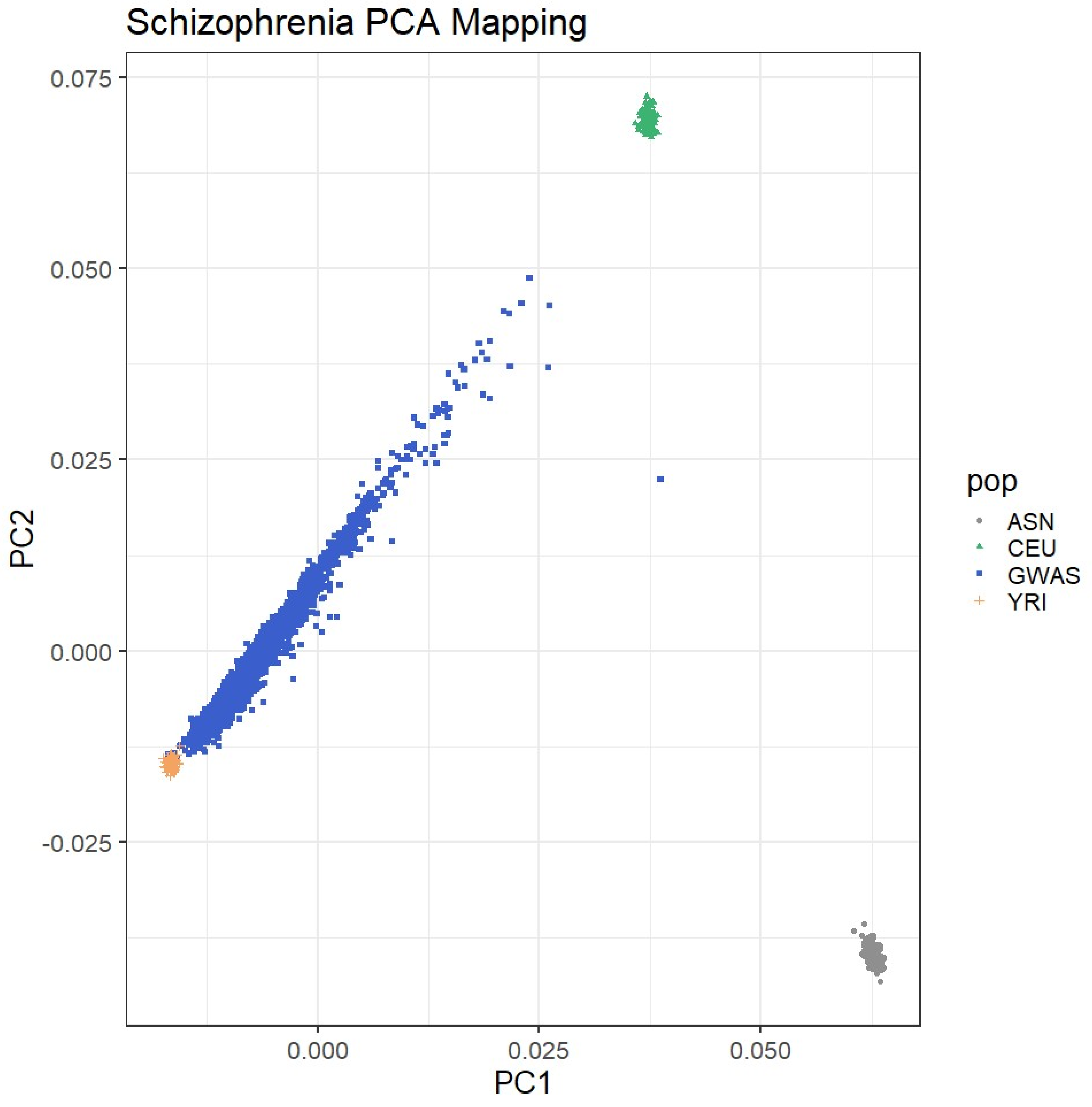

### Supplemental Figure 2

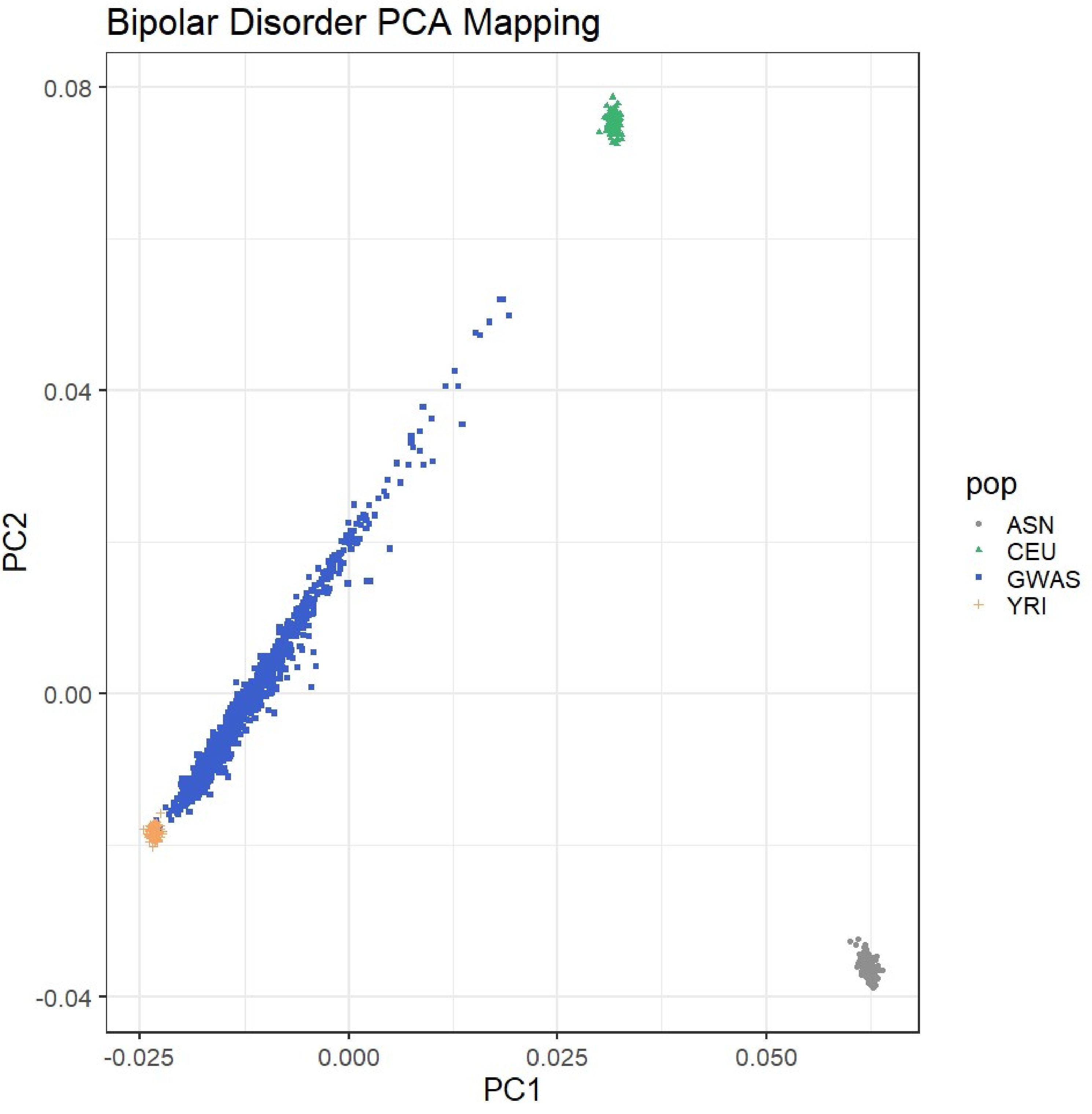

### Supplemental Figure 3

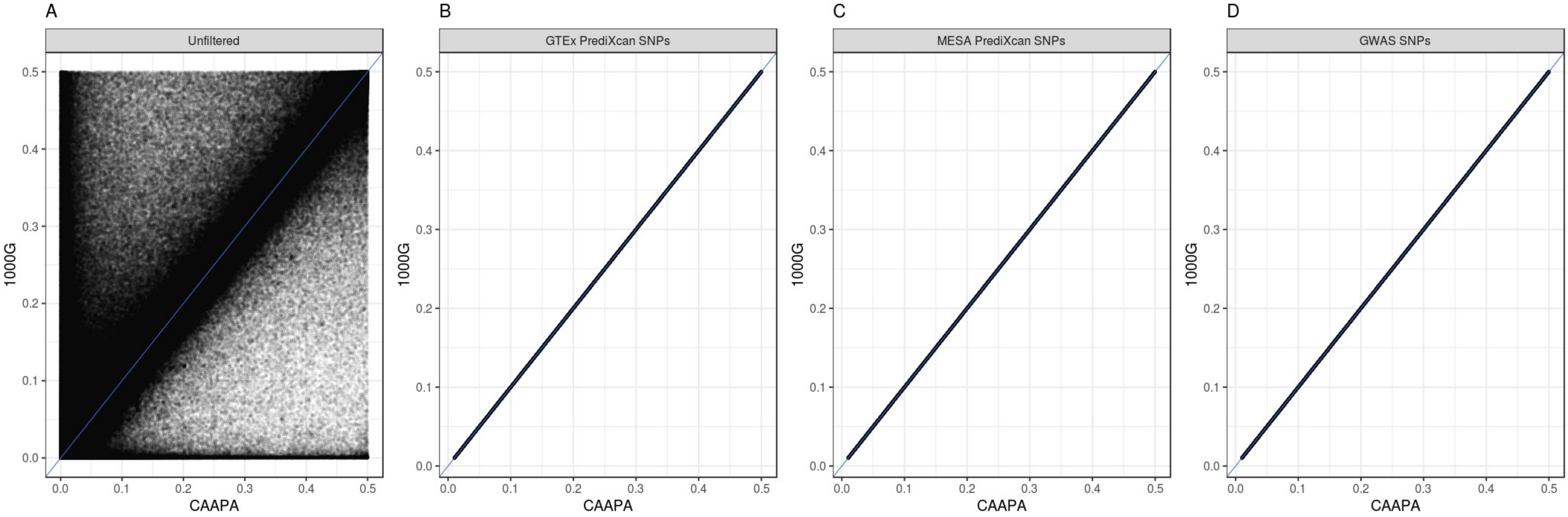
